# Biased estimates of phylogenetic branch lengths resulting from the discretised Gamma model of site rate heterogeneity

**DOI:** 10.1101/2024.08.01.606208

**Authors:** Luca Ferretti, Tanya Golubchik, Francesco Di Lauro, Mahan Ghafari, Julian Villabona-Arenas, Katherine E. Atkins, Christophe Fraser, Matthew Hall

## Abstract

A standard method in phylogenetic reconstruction for representing variation in substitution rates between sites in the genome is the discrete Gamma model (DGM). Relative rates are assumed to be distributed according to a discretised Gamma distribution, where the probabilities that a site is included in each class are equal. Here, we identify a serious bias in the branch lengths of reconstructed phylogenies when the DGM is used, with the magnitude of the effect varying with the number of sequences in the alignment. We demonstrate the existence of the bias in both simulated dataset and using real HIV-1 sequences; in both cases branch lengths are overestimated. We show that the alternative “FreeRate” model, which assumes no parametric distribution and allows the class probabilities to vary, is not subject to the issue. We further establish that the reason for the behaviour is the equal class probabilities, not the discretisation itself. We explore the mathematics of for the phenomenon, showing how maximum likelihood branch lengths under the DGM may differ from the true ones used to generate the tree, and that branch lengths will be overestimated where there is a long-tail of fast-evolving sites in the true rate distribution, which is the usual situation in real datasets. We recommend that the DGM be retired from general use. While FreeRate is an immediately available replacement, it is known to be difficult to fit, and thus there is scope for innovation in rate heterogeneity models.

## Introduction

It was recognised early in the development of molecular phylogenetics that it is unrealistic to assume that mutation rates do not differ between genomic sites (Fitch and Margoliash 1967; Uzzell and Corbin 1971). One key reason for the existence of variation is the degeneracy of the genetic code, where a random nucleotide substitution is least likely to be synonymous at the first codon position and most likely at the third, making mutations at the latter more likely to be neutral (Fitch 1986; Wolfe, Sharp, and Li 1989). In more general terms, the functional and structural role of sites in the genome vary very widely, and thus the evolutionary pressures on each, and hence the probability of observing a mutation there, cannot help but differ considerably (Yang 1996).

By the 1990s it was clear that methods were needed to account for between-site rate variation, and it became standard to use a gamma distribution to model it. An overall substitution rate *µ* is modified at each site in the genome by a draw from a gamma distribution with shape *α* and scale 1*/α*. (The mean of the distribution is thus 1 and the mean rate over the entire genome remains *µ*.) This was first employed in distance matrix calculations (Nei and Gojobori 1986), before Yang (1993) integrated it into maximum-likelihood algorithms. A year later, due to the computational limitations of research in the early 1990s, Yang (1994) introduced a pair of approximations. The first of these, the “discrete gamma model” (DGM), approximates the gamma distribution by dividing its support into a finite number (four was suggested) of contiguous regions of equal probability. The mean (or median) value of the gamma probability density function (PDF) on each region is calculated, and each site in the genome takes each of those values with equal probability. In subsequent years, this approximation became the absolutely standard approach to rate heterogeneity in almost all major software packages for maximum likelihood and Bayesian phylogenetic reconstruction, including PhyML (Guindon et al. 2010), RAxML (Stamatakis 2014), IQ-TREE (Minh et al. 2020), MrBayes (Ronquist et al. 2012), and both versions of BEAST (Baele et al. 2025; Bouckaert et al. 2019).

It might well be asked why, if one was to assume a discrete probability distribution for between-site rate variation, it needed to be assumed that either it took a gamma form or that every rate had an equal probability, and both assumptions can be relaxed. Yang (1995) outlined a general non-parametric approach, in which there are *k* rates *ρ*_1_, …, *ρ*_*k*_ and the probability that a site evolves according to each *ρ*_*i*_ is *w*_*i*_. Because ∑_*i*_ *w*_*i*_ = 1 and the mean relative rate across all sites is always assumed to be 1 (i.e. ∑_*i*_ *w*_*i*_*ρ*_*i*_ = 1) this model has 2(*k* − 1) free parameters. In contrast, the DGM has only one, the shape of the gamma distribution itself, regardless of how many categories are chosen. This increase in the number of parameters causes computational problems that continue to this day (Morel et al. 2021).

The DGM and FreeRate are both simple types of mixture model, and the use of mixture models for between-site rate variation has been a subject of continued development (Mayrose, Friedman, and Pupko 2005; Huelsenbeck and Suchard 2007; Evans and Sullivan 2012; Wu, Suchard, and Drummond 2013; Crotty et al. 2020). The version of Yang (1995) was belatedly christened “FreeRate” by Soubrier et al. (2012) and it is implemented under that name in the current versions of PhyML (Guindon et al. 2010), RAxML-NG (Kozlov et al. 2019), and IQ-TREE (Minh et al. 2020).

Previous critiques of the DGM have tended to demonstrate that, if the true rate distribution is not well-described by a gamma distribution, a more flexible model is superior for inference (Mayrose, Friedman, and Pupko 2005; Soubrier et al. 2012). Here we demonstrate a more fundamental problem that occurs even if true rates are actually drawn from a continuous gamma distribution, and which has profound implications for phylogenetic reconstruction. We demonstrate that branch lengths estimated using a DGM are biased and sometimes considerably so. The direction of the bias is towards overestimation for the simulated and real HIV datasets that we analysed. We further show that this effect increases with the number of sequences in the alignment. This phenomenon does not occur when FreeRate is used instead, and are a result of the equal size of the categories used in the DGM (rather than the discretisation itself or the choice of a gamma distribution). We demonstrate the problem using real and simulated data, and then provide an analytic demonstration of why it happens. We recommend, as a result, that the DGM be retired from use and that, if the issues with FreeRate render it unsuitable for general use, work is needed to produce a robust alternative.

## Materials and Methods

### Site rate heterogeneity models

Models of rate heterogeneity assume that each site *D*_*i*_ in the genome **D** is assigned a variable rate of evolution, *ρ*_*i*_ ∈ ℝ^+^. For the category of model we are dealing with, the *ρ*_*i*_ are assumed to be independent draws from a probability distribution *H*. This may be discrete or continuous. The rate at each *D*_*i*_ is then taken to be an independent draw from the distribution whose probability mass or density function is *f*_*H*_ .

This distribution is of a set of *relative* rates, and describes the heterogeneity among sites independently of the absolute rate of evolution. Thus its mean is taken to be 1, i.e.

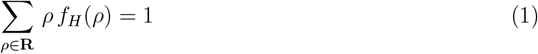

for a discrete *H* taking values in a finite set **R** of positive reals, or

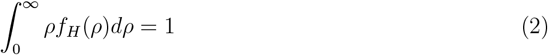

for a continuous *H*. When *H* is a gamma distribution, the requirement of a mean of 1 allows the distribution to be defined only by its shape *k*; the scale must then be 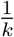.

A model of nucleotide substitution with a rate heterogeneity component has two random outcomes: the assignment of rates to sites, and the allelic states along the tree at each site. This latter, at the tips, produces the alignment, the data source in phylogenetic inference. The former are not directly observed. In the reconstruction models we are concerned with in this paper, which can be broadly classified as mixture models, their values are integrated over (Yang 1994). There is a separate class of “partition models”, which do perform per-site rate assignment as part of the algorithm, or allow the user to assign sites to categories prior to inference, but which we are not concerned with here (Lanfear et al. 2012).

Suppose *H* is discrete, the parameters of the rate heterogeneity model are *P*, and those of the nucleotide substitution model are *M* . The likelihood of a phylogeny **T** given a multiple sequence alignment **D** with *n*_sites_ positions is:

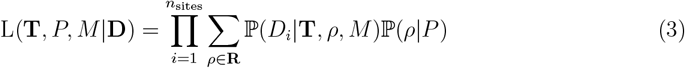

where *D*_*i*_ denotes the content of **D** at site *i*. Maximisation of this likelihood leads to maximum likelihood estimates for all parameters, denoted here by 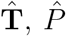, and 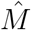 . Henceforth we refer to the terms P(*ρ*|*P*) as *weights*; they must sum to 1 over *ρ ∈* **R**.

The classic DGM belongs to a larger class of “equal-probability models”, where sites are assigned to one of *n*_cats_ rate categories where *n*_cats_ is finite. The important property here is that these categories are equiprobable, i.e. there is a fixed probability 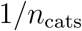 that a site belong to any one of them. Equivalently, in expression (3), the weights are identical.

Each category is assigned a specific relative rate. In the case of the DGM, the rates are chosen as either the mean or the median (Yang 1994) of the rates in the corresponding *n*_cats_-ile of a Gamma distribution. In the case of the mean, to which we give the full treatment here (the behaviour observed in this paper is unaffected by choosing the median instead) this is given by:

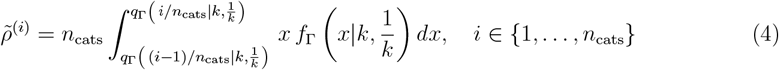

where *f*_Γ_ and *q*_Γ_ are respectively the PDF and quantile function of a Gamma distribution with shape *k* and scale 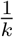. Note that the mean of the rates among categories is 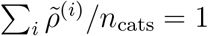 .

### Simulations

100 simulated phylogenies of 250 tips were created using the `rtree` function in the R `ape` package. For each, a real number *r* was sampled from the uniform distribution on the interval [-5, -2]. Branch lengths of the tree were rescaled such that their mean was 10^*r*^, simply by multiplying all simulated branch lengths by a constant value. Each of these trees was then used to generate sequences with AliSim (Ly-Trong et al. 2022), with 25,000 bases and the GTR rate matrix model with equal base frequencies. For a site heterogeneity model, we simulated using:

1. A continuous gamma distribution with that shape parameter of 0.5 and a scale parameter of 2.
2. A 4-category discretised version of the same gamma distribution, with each category taking up 1/4 of the probability space and its value on that quarter being the mean of the PDF in that interval.

Option 1 is *true* gamma-distributed rate heterogeneity, while in option 2 sequences are generated using the DGM itself. Phylogenies for each alignment were reconstructed using IQ-TREE 2.2.0 (Minh et al. 2020) specifying GTR+F+G4 (4-category DGM) and GTR+F+R4 (4-category FreeRate) as substitution models.

In a second exercise, we did this again, but this time fixed the mean branch length at 0.01 substitutions per site. Instead, what we varied was the number of tips in the simulated tree, between 50 and 500 in increments of 50. We did ten replicates of this process. We only simulated using option 1 in this case, but reconstructed using both the DGM and FreeRate as before. We also analysed these alignments in the same way using phyml version 3.3.20220408 (Guindon et al. 2010) and RAxML-NG version 1.2.2 (Kozlov et al. 2019).

### HIV phylogenetic reconstruction

For real data, we took an alignment of 1000 full-genome subtype C HIV sequences from the PopART Phylogenetics study (Hall et al. 2024; Hayes et al. 2019), available via the PANGEA-HIV consortium (Abeler-Döorner et al. 2019). Sequences can be downloaded at http://github.com/PANGEA-HIV/PANGEA-Sequences. Positions where more than 10% of sequences had an N or gap character were removed, and the whole alignment consists of 7854 base pairs. This alignment was progressively downsampled by subtracting 25 random sequences from it, to give a total of 39 alignments of between 50 and 1000 sequences; the 50 in the former are shared between all of them. Reconstruction was performed using GTR+F+G4 and GTR+F+R4 as above.

As a sensitivity analysis, we also performed this exercise using 6-category, 8-category and 10-category DGMs (GTR+F+G6, GTR+F+G8 and GTR+F+G10) as well as the 4-category DGM with invariant sites (GTR+F+I+G4), using the same set of alignments as above, and also repeating it with three other alignments of subtype C PANGEA sequences.

As a separate exercise, we used ModelFinder (Kalyaanamoorthy et al. 2017) to determine the preferred substitution model for each of the downsampled alignments.

## Results

### A bias in reconstructed branch lengths exists for the DGM but is absent for FreeRate

In figure 1a, we plot the mean branch length in the reconstructed phylogeny against the true mean branch lengths in the simulated phylogeny, by simulation method (true, continuous gamma distribution or discretised gamma distribution) and reconstruction method (DGM or FreeRate). For all but one combination, the reconstruction is near-perfect and shows no particular bias. The exception is where the generating distribution was the continuous distribution and the reconstruction the DGM, where branch lengths are overestimated, with the effect increasing with the mean true branch length. In figure 1b, we plot the true branch length against the ratio of reconstructed and true branch lengths, for the biased combination only. There is no appreciable bias for true mean branch lengths up to around 0.0005 substitutions per site, but after that the magnitude of the effect increases rapidly.

**Figure 1:**
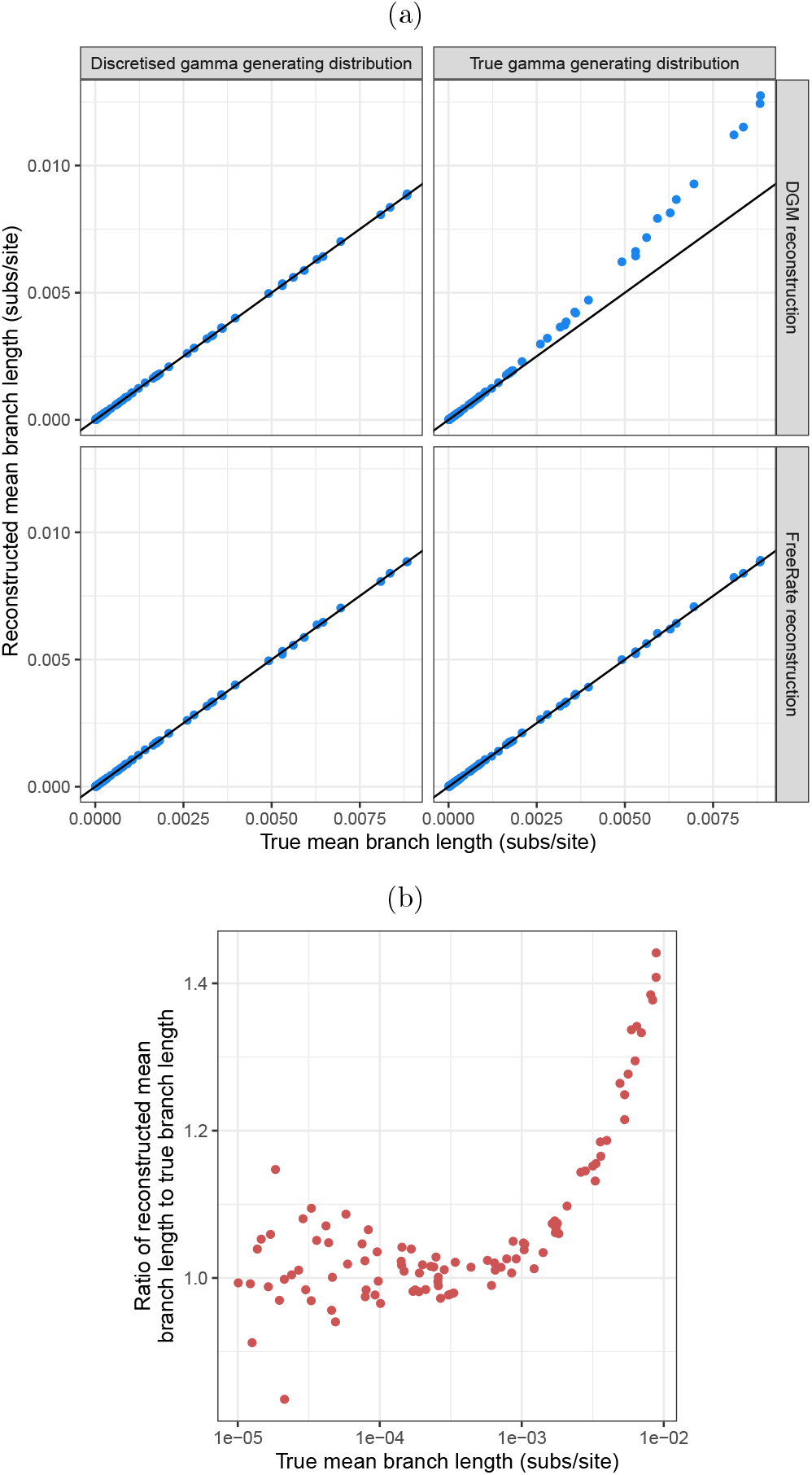
Comparison of reconstructed mean branch lengths to true branch lengths in simulated phylogenies. In a), we plot true mean branch length against that in the reconstruction. This is shown for generating gamma distributions that are either the true continuous distribution or the discretised version (that used by the DGM), and for the DGM and FreeRate as the reconstruction method. The black line is the line of equality. In b) for the continuous/DGM combination only, we plot the ratio of reconstructed to true mean branch length against true mean. Note that the x-axis in b) is on a log scale.

### The magnitude of the bias varies with the size of the alignment

In figure 2, the mean branch length in the generating tree is fixed to 0.01 substitutions per site, and the number of tips in that tree is varied instead. It can be seen that not only are DGM reconstructions biased, but the magnitude of that bias is highly dependent on the number of tips in the tree. For FreeRate, there is a slight suggestion of overestimation for small trees, but the effect is of nowhere close to the same magnitude.

**Figure 2:**
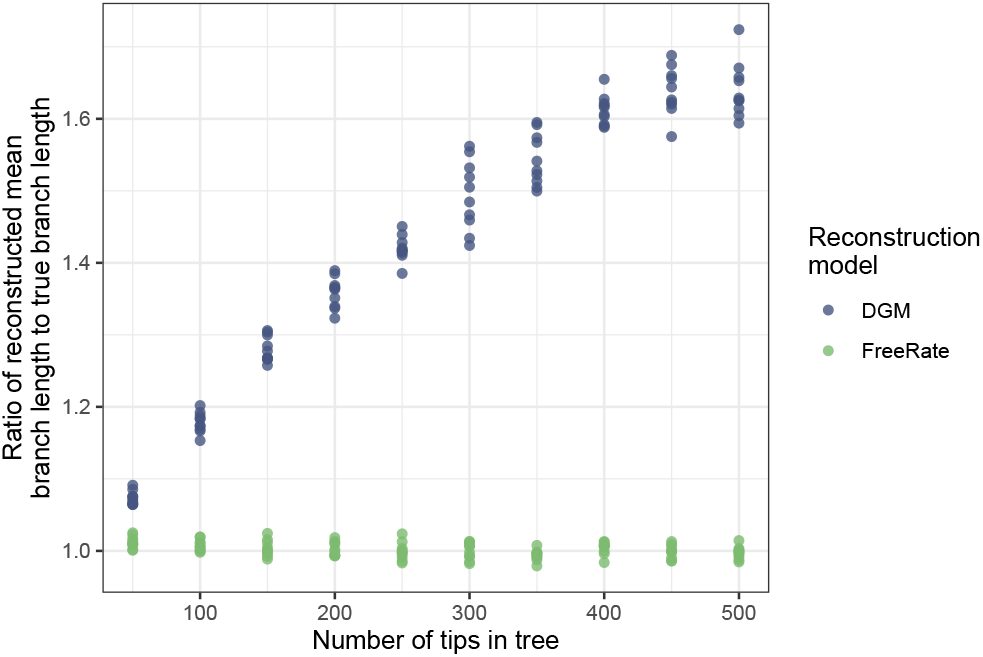
The effect of the number of tips in the tree. For a total of 100 simulated trees with varying numbers of taxa, the ratio of the reconstructed mean branch length to the true mean branch length (which was always 0.01 substitutions per site) is plotted. The generating rate heterogeneity distribution was a continuous gamma distribution; the reconstruction was done with the DGM (blue) and FreeRate (green).

While the main text results were obtained using IQ-TREE, figure S1 presents the equivalent of figure 2 for both PhyML and RAxML-NG. There is no meaningful difference in the output.

### Branch lengths in HIV phylogenies increase with sample size with the DGM, but not FreeRate

In figure 3, we plot, for the downsampled alignments up to size 250, patristic distances between the tips representing the 50 sequences common to all our HIV alignments. The comparison is between the reconstruction using just those sequences (*x*-axis) and in that using larger alignments (*y*-axis). While we have no ground truth for real data, the inflation of patristic distances as sample size increases when the reconstruction is done with the DGM is obvious and mirrors that presented in the previous section. The phenomenon is absent for FreeRate.

**Figure 3:**
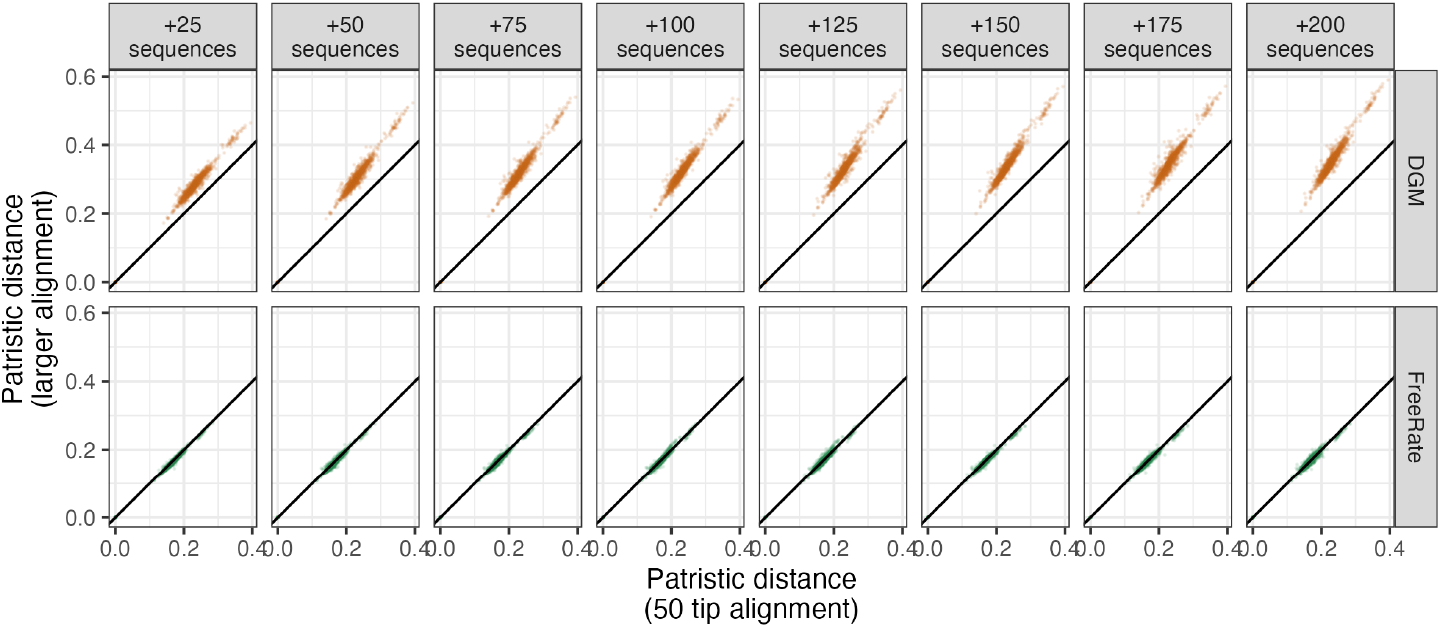
Patristic distances between the tips representing 50 subtype C HIV sequences in phylogenetic reconstructions. The x-axis gives these when the alignment consists of just those 50 sequences, while the y-axis are the same from a larger alignment, with the total number of sequences increasing in the facets from left to right. Top row: reconstruction uses 4-category DGM. Bottom row: reconstruction uses 4-category FreeRate.

Figure 3 suggests a linear relationship between patristic distances in datasets of different sizes, and so figure 4 displays the slope of regression lines for these comparisons, up to a total of 1000 tips. A rapid increase in the slope when moving from 50 to 125 tips indicates a swift increase in the amount by which branch lengths are overestimated. The effect does appear to eventually saturate after 250 tips, at which point there is little sign of a trend.

**Figure 4:**
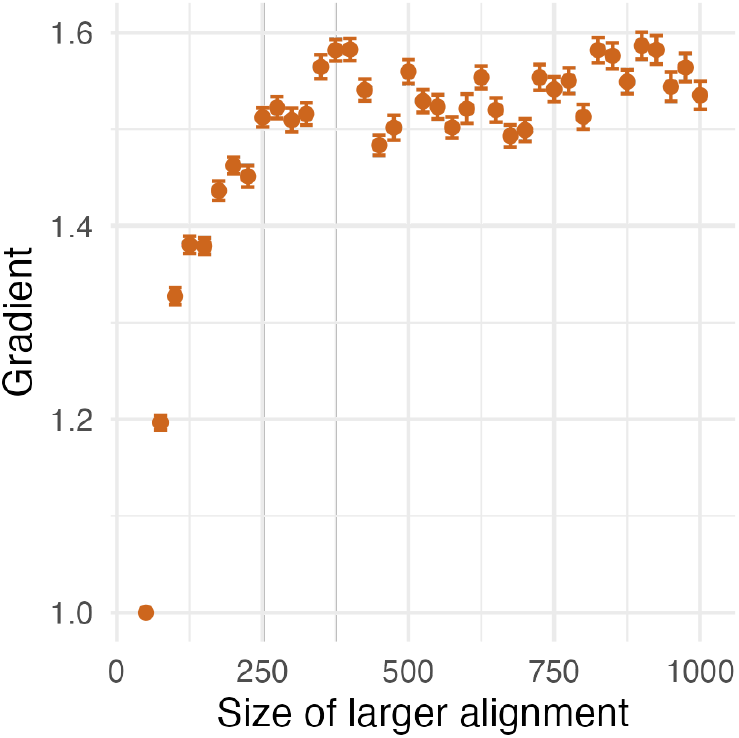
The slope of a linear regression line plotting patristic distances in phylogenies generated using the DGM on a 50-sequence alignment and in phylogenies from larger alignments. The x-axis is the size of the larger alignment. The error bars give the 95% confidence interval of the regression coefficient. In the 50-tip case the tree on both axes is the same one.

When ModelFinder was applied to these HIV alignments, the preferred model by Bayesian Information Criterion score was GTR+F+I+R10 (10-category FreeRate with an invariant sites category) for all sample sizes of 175 or above. For smaller alignments the invariant sites category could be omitted or the number of FreeRate categories decreased, but the DGM was never preferred.

If we increase the number of rate categories in the DGM, the issue is actually exacerbated. In Fig. 5, we plot slopes for models with a varying number of categories for four different 1000-sequence alignments; replicate 1 is that of figure 4. For a comparison of individual patristic distances from replicate 1, see figure S2. A 6-category DGM results in considerably more branch length expansion than a 4-category one. 8- and 10-category versions show more expansion still, although the progression is not straightforward and the 8-category version is often worse than the 10-category. On the other hand, adding an invariant sites category does mitigate the issue. This effect is, however, slight and the problem remains in evidence. Also of note is the considerable inconsistency in the behaviour of the expansion between replicates for larger number of categories; while the 4-category DGM converges to a slope of around 1.5 in all four replicates (and whether or not an invariant sites category is present), this kind of consistent behaviour is not in evidence for more complex models. In some cases (most clearly replicate 2 for the 6-category DGM), the largest alignments show less expansion than some of the smaller ones. Figure S2 suggests that the scaling effect on branch lengths can be somewhat less linear for the more complex models.

**Figure 5:**
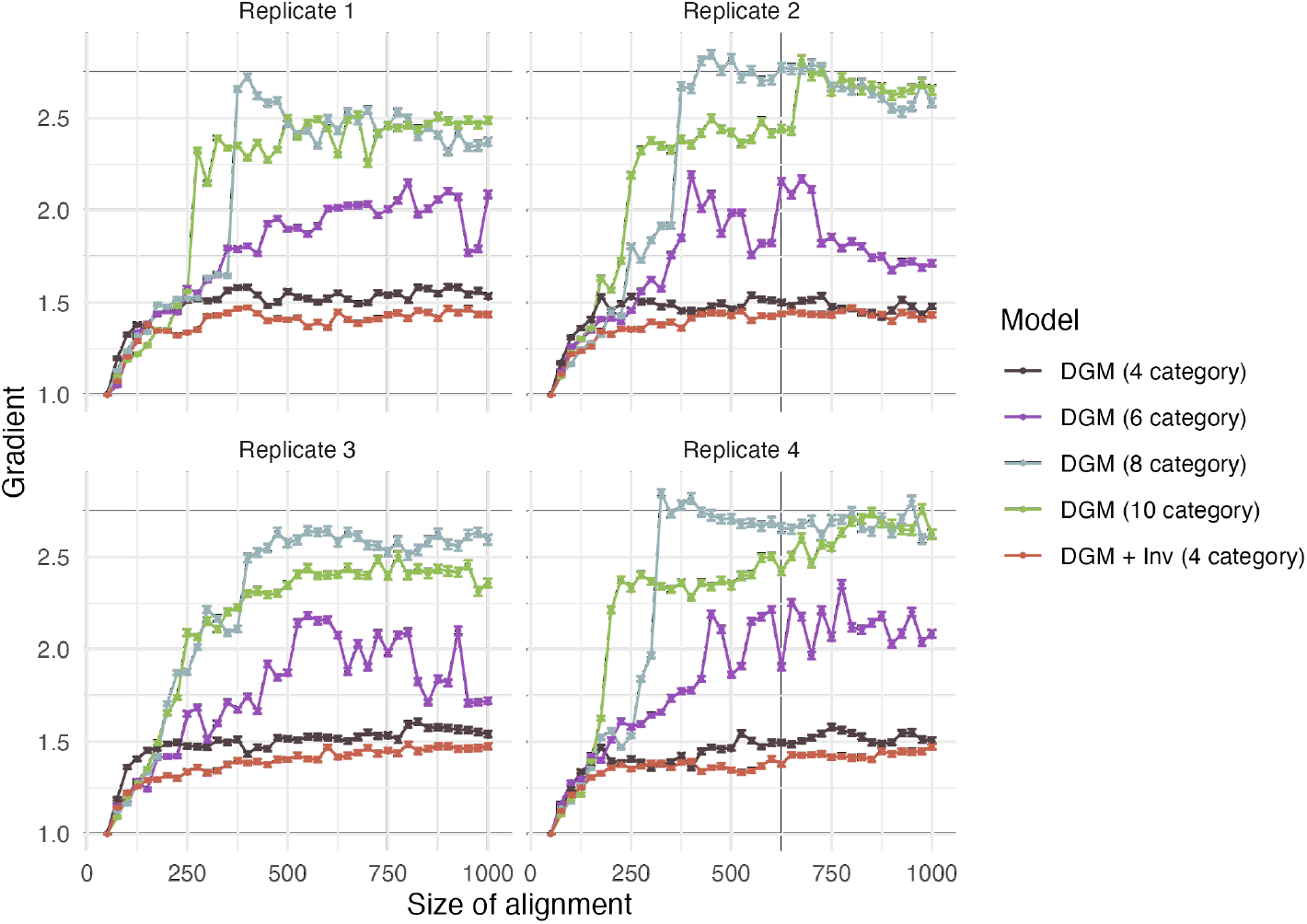
The slope of a linear regression line plotting patristic distances in phylogenies generated using the DGM on a 50-sequence alignment and in phylogenies from larger alignments. The x-axis is the size of the larger alignment. The error bars give the 95% confidence interval of the regression coefficient. In the 50-tip case the tree on both axes is the same one. Points are coloured by the version of the DGM used for reconstruction. The four facets correspond to different initial 1000-sequence alignments which were progressively downsampled, all consisting of subtype C HIV sequences from PANGEA-HIV.

### Biased branch length estimates as the result of equiprobable rate categories

To summarise the previous results, the DGM gives good estimates of branch lengths when the generating model is itself discretised, but an obvious bias occurs when attempting to fit it to sequences generated using a truly continuous gamma model. This is most pronounced for trees with longer branches, and its magnitude increases with the number of sequences in the alignment.

These simulations notably lack any process mimicking misalignment, recombination, sequencing error, selection, or any other complex biological processes that might drive the phenomenon. As none of these are present, they cannot explain the results. The bias is due simply to the generation of a model using a continuous distribution and reconstruction using the DGM. As FreeRate branch lengths are largely unbiased, it would seem natural that it is also correctly estimating the form of the input distribution even if its nature as a gamma distribution is not specified, and this, indeed, is the case. In figure 6, we plot, for four example alignments of 500 sequences from the simulations of section “The magnitude of the bias varies with the size of the alignment” above, the following quantile plots:

- The true, continuous quantile plot of the generating gamma distribution (black).
- The quantiles of the fitted FreeRate distribution (purple).
- The quantiles of the true gamma distribution discretised according to the FreeRate weights (teal).
- The quantiles of the fitted DGM (green).
- The quantiles of the true gamma distribution discretised with equal weights (red); this is independent of the alignment and identical on each plot.

**Figure 6:**
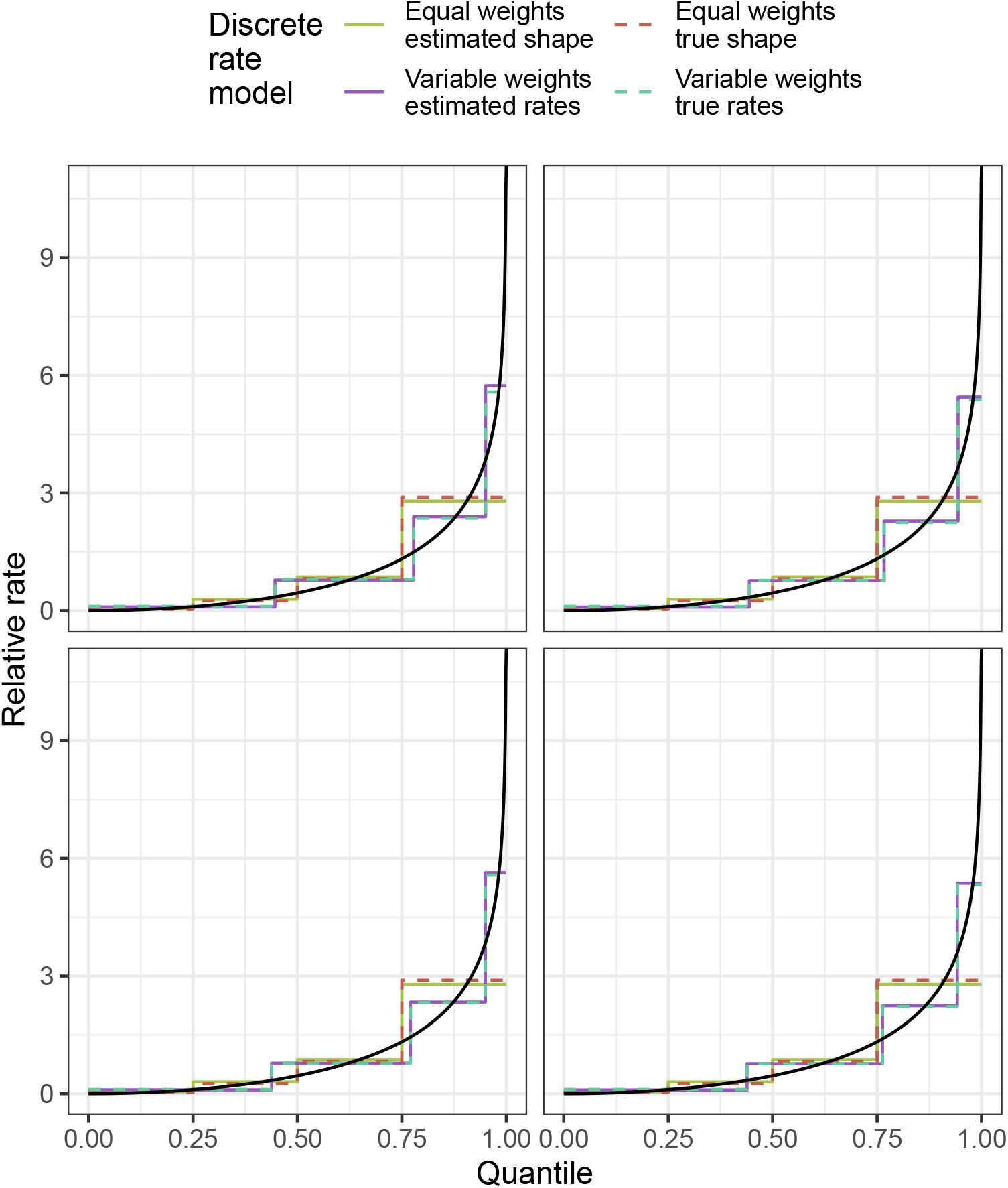
For four simulated alignments of 500 sequences each, the actual quantile plot of a gamma distribution with shape parameter 0.5 and scale parameter 2 (black), and of the DGM discretisation of that distribution into four categories (red). These are compared tothe quantiles of the fitted FreeRate distribution (purple), the quantiles of the fitted DGM (green) and the mean values of the true gamma distribution on the quantiles defined by the fitted FreeRate weights (teal).

The FreeRate reconstruction is not subject to the branch expansion problem. The teal line represents a particular discretisation of the true rate distribution, and the purple one the estimated distribution according to FreeRate, which is unaware that the true distribution is a gamma. These lines follow each other very closely, which suggests that FreeRate can estimate a discretised Gamma distribution in such a way that branch length estimates are not biased. No standard maximum likelihood phylogenetics package currently allows the user to specify, as a rate heterogeneity model, a discretised gamma distribution whose weights are variable and estimated by the algorithm. However, it would naturally follow that if FreeRate is able to mimic the form of one without being subject to branch length bias in the output, such a “gamma + variable weights” option in a package would also be able to. The problem thus cannot be the discretisation, and must instead lie with the equal weights.

We conclude that reconstruction of phylogenies using the DGM from alignments whose true rate heterogeneity model is a continuous gamma distribution are systematically biased, and that the reason is the constraint that the probabilities of a site belonging to each category are equal.

### Branch expansion is a consequence of model mis-specification and a tail of rapidly evolving sites

In the appendix to this paper, we delve further into the mathematics of the phenomenon. We summarise those results here. Under “Likelihood expression for star-like phylogenies” we give an expression for the likelihood of branch lengths in a star-like tree (a tree with a single internal node), after some simplifying assumptions. Maximising this expression allows us to estimate the magnitude of the bias in branch lengthestimates, for various scenarios of true branch length and alignment size (figure 7). We see that a longer branch length is initially associated with a faster increase in maximum likelihood branch length as the number of tree tips increases, but that there is also an asymptotic limit to the bias which is obtained sooner, and at a lower value.

**Figure 7:**
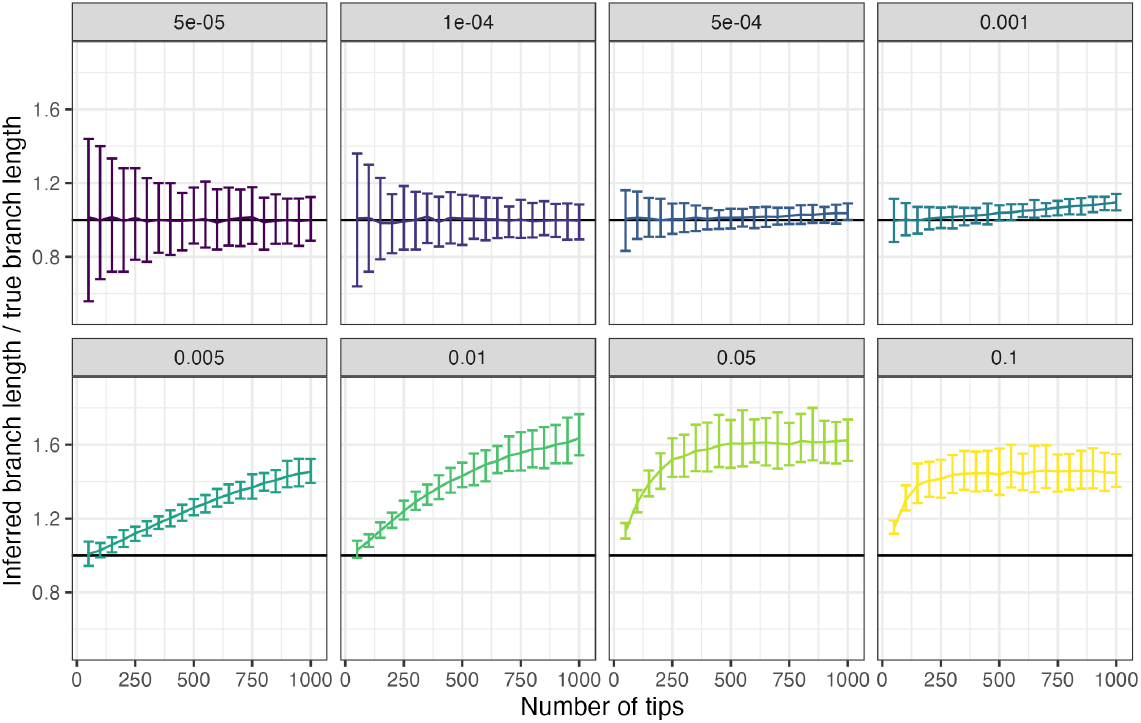
Ratios of the maximum likelihood estimate of the (constant) phylogenetic branch length in a star-like phylogeny to its true length, for different sizes of alignment (x-axis) and lengths of the true branch (substitutions per site, facets). Calculations are the result of simulating 100 alignments of 5,000 sites each with a gamma shape parameter of 0.5. The central line connects the mean maximum likelihood estimate across those 100 alignments, with the error bars giving the 95% symmetric quantile around the median.

That the DGM is a misspecified model of the true distribution of rates would seem clear for real data, as it was always intended as a simplified method (Yang 1994). However, the bias exists even if the generating rate distribution is a continuous gamma distribution, and, as shown above (“Branch lengths in HIV phylogenies increase with sample size with the DGM, but not FreeRate”) it clearly also occurs in genuine datasets.

In the section entitled “And equal-weights model misapplied to a discrete generating model with unequal rates” we give an expression for the bias in branch lengths when a discrete equal-weight distribution is assumed but the true model of evolution follows a FreeRate distribution with the same number of categories but variable weights. We show that branch lengths are overestimated when there is a negative correlation between the weight of each category *w* and its relative rate *ρ* in the true model of evolution. More precisely, if the true branch length is *λ* and the estimate 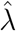, and **P** and **W** are the vectors of rates and weights respectively, the relative bias is given by:

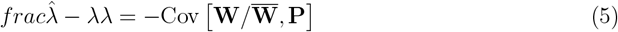

If rates and weights are positively correlated, i.e. larger rates are most common, this expression is negative and branch lengths are underestimated. However, this is likely be rare in the real world as most inferred rate distributions have a mode of less than 1 (Jia, Lo, and Ho 2014; Silvestro, Latrille, and Salamin 2024). It would be much more common to see a negative correlation between rate and weight, leading to overestimation of branch lengths.

Finally, we show analytically that the overestimation of branch lengths also occurs in a scenario where the generating distribution is continuous. If the generating model is a continuous Gamma distribution with shape and scale equal to 1, but the phylogenetic reconstruction is performed using a discrete model with two categories of equal rates, it can be shown analytically that the branch lengths are overestimated by about 20%.

## Discussion

This is, to our knowledge, the first time this bias has been described in the literature, despite the almost thirty-year history of the DGM. Brown et al. (2010) attributed a similar phenomenon, observed in the Bayesian realm only, to either poor Markov Chain Monte Carlo mixing or excessive posterior weight to trees with long branches. On the maximum likelihood side, Nguyen, Haeseler, and Minh (2018) found that problematic heuristic optimisation routines in the major packages were responsible for misestimation of the shape parameter of the DGM and tree length, as well as the proportion of invariant sites. Mannino et al. (2020) identified a different problem with the equal-probabilities specification, in that it places a hard upper bound on the variance of the rate distribution, but did not explore branch lengths. Schwartz and Mueller (2010) explored bias in branch lengths in relatively small trees using both Bayesian and maximum likelihood methods. They observed a tendency for underestimation in both cases, along with some overestimation of longer branches for maximum likelihood. Their focus was in comparing estimates for individual branches to their true lengths, rather than comparing reconstruction results as we do here, and while they did employ the DGM, the datasets used may have been too small (in terms of numbers of taxa) for the phenomenon we observe here to have become obvious. The phenomenon we identify here is broader than all of these, with major implications for the field.

The equal weights specification is a highly problematic choice, which causes systematic bias in the estimation of phylogenetic branch lengths. In our wider work this has almost always been in the direction of overestimating them. This effect increases with sample size but can still be significant for small alignments if branch lengths are long enough. It should be emphasised that nothing about this phenomenon is biological. It is not specific to any organism, although its magnitude will depend on quantities such as the branch lengths in a typical phylogeny. It is not even specific to biological organisms at all; studies using phylogenetic reconstruction in the realm of linguistics have also used the DGM in the past (e.g. Lee and Hasegawa (2011)). The effect is more pronounced when mutation rates are fast (Fig. 2), and as such it is perhaps not surprising that we first observed it when analysing the genomes of RNA viruses. That the simulations here did not show the bias when mean branch lengths were less than around 0.0005 substitutions per site should be not be taken to indicate that it is completely absent below that. An implication of figure 7 is that smaller branch lengths lead to a slower emergence of the phenomenon as sample size increases, but it does eventually emerge for large alignments. We were not able to analytically establish the reasons for this variation in the rate of emergence in this work. The vast diversity of the types of data and sizes of datasets used in phylogenetics were impossible to fully capture in our simulation work here. As a result, we would recommend that researchers check using FreeRate or a similar model that their data is not subject to the issue, even if their branch lengths would seem sufficiently short to have avoided it.

An obvious line of further enquiry, which we do not explore here but is the subject of intensive ongoing work, is the effect of this bias on time tree inference (Sagulenko, Puller, and Neher 2018; Baele et al. 2025; Bouckaert et al. 2019). In a time tree, a molecular clock model is employed so that branch lengths are in units of calendar time and tips appear at points on the timeline corresponding to the dates upon which the corresponding samples were taken. Longer branches in the underlying divergence tree will not necessarily correspond to longer branches in a time tree, because a model can instead increase the rate of evolution dictated by the molecular clock. We can compensate for the allocation of a greater amount of evolution to a particular phylogenetic branch by, in turn, increasing our estimate for how quickly that evolution happened. However, time tree models are often complex, and the extent to which the effects of this bias affect branch lengths and the extent to which they affect clock rates requires urgent investigation.

Increasing the number of categories in the DGM does not resolve the problem and in fact makes it worse (Figs. 5, S2). On the other hand, adding an invariant sites category appears to be slightly mitigating, at least for our HIV alignments. This is perhaps because the size of that category is estimated and as such the model does, in a limited way, enable variable weights. Nevertheless, the reduction is modest and we would not recommend it as a long-term solution. For six or more categories, the behaviour of the expansion can be strikingly different when applied to different alignments. Identifying the reasons for this is beyond the scope of this work, but we would suggest that limitations of the heuristic explorations of tree space would need to be excluded as a cause before a mathematical explanation was sought.

FreeRate is the most immediate option for an alternative to the DGM that is not subject to the problem of branch length bias. It is implemented and documented in state of the art maximum likelihood phylogenetics packages (Stamatakis 2014; Minh et al. 2020) although much less common in Bayesian packages. For HIV, we always found that ModelFinder preferred it to the DGM. It is, however, known to be difficult to fit in a maximum likelihood framework (Morel et al. 2021). We stress that our objective in this paper is to inform the research community of a particular issue with the DGM, not to advocate for FreeRate as a gold standard alternative. A potential alternative from the existing literature, proposed by Felsenstein (2001) is to use generalised Gauss-Laguerre quadrature to approximate the integral of the likelihood over the rate categories. We would also suggest that the time is right for innovations in site rate heterogeneity models, after so many years during which the DGM has been used by default. This process may be underway, as examples taking a machine learning approach are now starting to emerge (Silvestro, Latrille, and Salamin 2024).

In summary, we would argue the problem that we identify here regarding the DGM is one that should preclude its future use. With our HIV data we showed that branch lengths are highly influenced by the size of the alignment, a situation that cannot be compatible with robust inference. FreeRate provides one option, but has its own disadvantages. The impact of this problem on phylogenetic dating is less clear and work is needed to establish the extent to which estimates may be changed in the light of this finding.

## Funding Acknowledgment

The computational aspects of this research were supported by the Wellcome Trust Core Award Grant Number 203141/Z/16/Z and the NIHR Oxford BRC. The views expressed are those of the author(s) and not necessarily those of the NHS, the NIHR or the Department of Health.

This study was supported by the Bill & Melinda Gates Foundation through the PANGEA consortium (OPP1175094 and OPP1084362) and by the Wellcome Trust through the ARTIC Network Collaboration Award (206298/Z/17/Z).

CJVA and KEA were funded by an ERC Starting Grant (award number 757688) awarded to KEA.

TG is supported by an Investigator Grant (GNT2025445) from the National Health and Medical Research Council, Australia (NHMRC).

## Acknowledgment

We thank Sergei Kosakovsky Pond and Andrew Rambaut for helpful suggestions.

## Data availability statement

No new real data were generated or analysed for this research. A DRYAD repository for this paper is available at https://doi.org/10.5061/dryad.d51c5b0fh. This contains all simulated alignments. The real HIV sequence data used are available at https://github.com/PANGEA-HIV/PANGEA-Sequences, and the DRYAD repository lists all sequences used in the four replicates.

Code to perform the analysis of appendix section “Numerical results” can be found at https://zenodo.org/records/15276300.

## Appendix

### Summary

In this appendix, we:

1. provide a brief reminder of the mathematical concepts involved in continuous time Markov chain (CTMC) models of character evolution (“Back to basics”)
2. introduce a simplified phylogenetic scenario in which we will work (“Simplifying assumptions”)
3. derive an expression for the likelihood of branch lengths in this scenario (“Likelihood expression for star-like phylogenies”)
4. show the existence of the bias by numerical estimation of the maximised likelihood (“Numerical results”)
5. give a general argument, for any tree, as to why a mis-specified model may lead to biased estimates of branch lengths, and show that this applies in a simulation exercise (“Biased per-site rate estimates are accompanied by biased branch lengths”)
6. in our simplified scenario, show that in the limit of a large number of tips, and of sufficiently small branch lengths, we would not expect estimates under an equal-weights model to match true branch lengths (“Likelihood in the large *n*_tips_ limit”)
7. show that, in a scenario where an equal-weights model is used to reconstruct a phylogeny when the true underlying distribution is discrete but does not have equal weights, there is a biasing effect which will usually be in the direction of overestimation of branch lengths (“And equal-weights model misapplied to a discrete generating model with unequal rates”)
8. show that the effect is also present when the true underlying distribution is a continuous Gamma distribution (with shape and scale equal to 1) and the reconstruction model is a two-category discrete equal-weights model (“An equal-weights discrete model as an approximation for a continuous Gamma generating model”)
9. demonstrate how the effect persists when some of the key assumptions in our simplified can be relaxed (“Generalisations”)
10. finally, acknowledge that our simplifying assumptions leave some questions about the issue unresolved (“Open questions”)

### Back to basics

Statistical phylogenetic inference, whether maximum likelihood or Bayesian, has as its core the likelihood function (Felsenstein 1973):

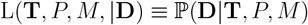

where **T** is the tree, **D** the sequence alignment (that is, the data), *P* the parameters of the rate heterogeneity model and *M* the parameters of the nucleotide substitution model. Implicit in this formulation is that **T** exists independently of **D**; there is an underlying “true” tree, with branch lengths in substitutions per site, even if no set of sites is specified. (This is perhaps most easily recognised in the scenario of generating simulated sequences, as packages generally take a fixed tree as input and then provide an alignment of any desired length.)

A component of *M* is an overall rate of evolution *µ*, such that the expected number of substitutions at a single position during an “evolutionary time period” of length *t* is *µt*. As the units of “evolutionary time” are arbitrary, one can actually assume *µ* = 1, as is common in phylogenetics packages.

**D** must have a finite length; suppose this is *n*_sites_. *P* is a vector of positive reals 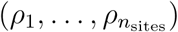 providing a (not necessarily unique) scale factor for *µ* at each site *D*_*i*_ of **D**. The likelihood is then calculated as:

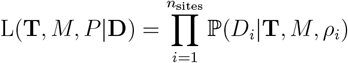

If *µ* is to remain equal to 1, then we need 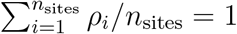.

The exact assignment of rates to sites is, of course, unknown outside of a simulation context (and indeed the entire concept of reasonably simple parametric distribution of rates is a considerable simplification of the mutational process for real data). In our setting of a finite mixture model scenario the possible values of the *ρ*_*i*_ are members of a finite set **R** and assignments are integrated over:

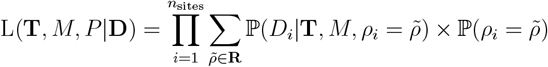

To write down an exact analytical expression for this likelihood requires a large number of choices regarding how the mutational process is assumed to behave. For example, one must choose how many of the relative rates of different types of base substitution are taken to be the same (Darriba et al. 2020), and the exact nature of **R**.

Maximum likelihood phylegenetics provides an estimate 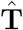 for **T**, as well as 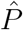 for *P* and 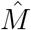 for *M* .

### Simplifying assumptions

Let us simplify **T** considerably by assuming the following:

1. **T** is star-like; all tips are direct children of the root, which is a polytomy of order *n*_tips_.
2. All branches of **T** have the same length *λ*. The total length of the branches of **T** is *n*_tips_*λ*.
3. The alignment **D** consists of *n*_sites_ sites which take only two allelic states (0 and 1). The substitution model, before rate heterogeneity is considered, is simply transition from 0 to 1 or vice versa at an instantaneous rate of 1, and nucleotide frequencies are equal. For each site *i*, a rate heterogeneity parameter *ρ*_*i*_ is drawn from an unspecified probability distribution *H* with support on the positive reals and a mean of 1.
4. The root state of the phylogeny is known to be 0 at every site.
5. The topology of **T** and the form and parameters of the substitution model and rate heterogeneity model are known and need not be estimated.

As the expected number of substitutions at site *i* along a branch of length *λ* is *λρ*_*i*_, the expected number of substitutions across all sites is ∑ _*i*_ *λρ*_*i*_ and the expected number of substitutions per site is 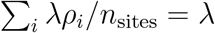

### Likelihood expression for star-like phylogenies

We want to obtain an estimate 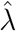 for *λ* using the observed alignment **D**. Let *K*_*i*_ *≤ n*_tips_ be the number of sequences with an observed substitution (i.e. a 1) at the *i*th site. As the overall substitution rate *µ* is 1, the substitution rate at site *i* is just *ρ*_*i*_. The *Q* matrix of the CTMC at that site is 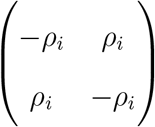, which has eigenvectors 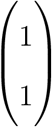 and 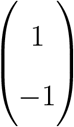for its eigenvalues 0 and −2*ρ*_*i*_. The matrix of eigenvectors is 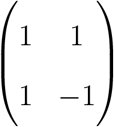 with inverse 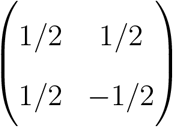 . Then

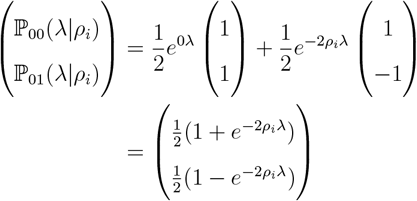

is the vector of probabilities for going from the state 0 to 0 or to 1 along a branch of length *λ*.

Suppose we are to estimate *λ* by maximum likelihood, but using a discretised rate heteregeneity distribution 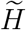with equal weight 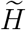 has *n*_cats_ rate categories with a set of relative rates 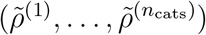 such that 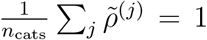 . The expected number of substitutions at site *i* along a branch of length *κ*, if *i* belongs to the category with rate 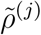, is 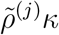. Thus if 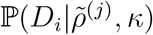is the probability of observing the data at site *i* if all branch lengths are *κ*,

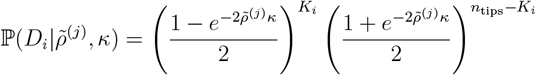

If 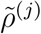is unknown then it needs to be summed over:

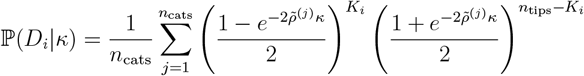

and then the probability of the entire alignment is:

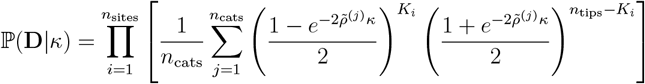

Treated as a likelihood (i.e. a function of *κ*), and taking logs, we get an estimated branch length 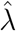:

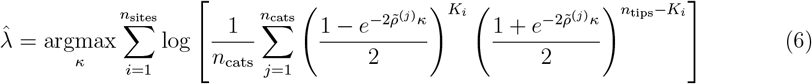

### Numerical results

We can calculate 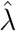 using (6), for a known *n*_sites_ and set of 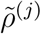, using standard numerical optimisation. We simulate evolution along *n*_tips_ branches of length *λ* and calculate *K*_*i*_ by counting the number of 1s at the end. Figure 7 plots maximum-likelihood estimates of 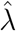 inferred using the 4-category Gamma model with alpha=0.5, against the actual branch length *λ*, for an alignment of 5,000 sites and a true gamma shape parameter of 0.5.

For small trees and slow rates, no bias in branch lengths is in evidence. The branch expansion effect is plain for longer branches, and the rate at which it occurs is correlated with *λ*. However, this increase eventually converges to an asymptote which is also reached faster for longer branch lengths, with the asymptotic value actually declining as the branch length increases. The latter effect may well be due to mutational saturation, as the increasing prevalence of unobserved back-mutations will tend to decrease the branch lengths in the reconstruction.

### Biased per-site rate estimates are accompanied by biased branch lengths

Here we present a general argument that applies to any tree **T**. We denote by *λ*_**T**_ the total length of the tree in units of expected substitutions per site, i.e. the sum of all its branch lengths. *λ*_**T**_ is the length of the underlying phylogeny, before any alignment is generated or any package is used for reconstruction. The branch lengths correspond to the number of substitutions per site observed in a sequence where site rates are distributed as the actual site-rate distribution *f*_*H*_ . On the other hand, if site-specific rates for each position were to be inferred, we can calculate an inferred total length *κ*_**T**_ from the expected number of substitutions at each site.

The effect of this is to scale the theoretical branch lengths by the mean rate of evolution 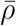 amongst the *n*_sites_ sites of the sequence:

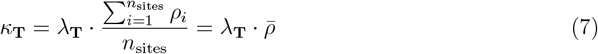

If all is well 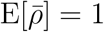, and the expected value of *κ*_**T**_ is *λ*_**T**_. However, this is not automatic once the expected value is conditioned on the data. Noting that (7) implies 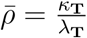, we have that

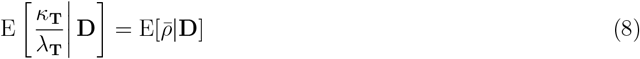

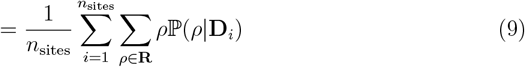

and, after applying Bayes’ Theorem to ℙ(*ρ*|**D**_*i*_), this is equal to:

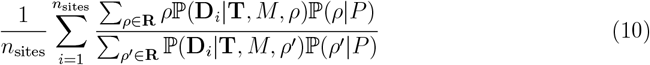

This is the mean of the empirical Bayes estimates of site rates across the whole genome (see equation 3 of Mayrose et al. 2004). There is no reason that it is necessarily 1.

From the point of view of inference, an ML phylogenetic algorithm generates an estimate *κ*_**T**_ for *λ*_**T**_. If the model misspecification causes the site-specific rates to be biased on average, i.e. 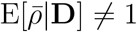, then *κ*_**T**_ will differ from *λ*_**T**_ by a multiplicative factor. Moreover, all individual branches of the tree will tend to be shorter or longer than the ones inferred by the model by the same factor.

ML software packages estimates of **T**, *M* and *P* can be used to estimate the expected substitution rate E[*ρ*_*i*_|**D**] at every site *D*_*i*_ in **D** using a variety of methods, one of which is empirical Bayes (Mayrose et al. 2004). We can use this to confirm using simulated data that the ratio 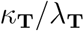is approximately equal to the mean expected substitution rate 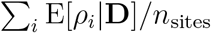.

A simulated HIV transmission tree was generated using PopART-IBM (Pickles et al. 2021). The result was a completely sampled epidemic of 1771 cases. This was transformed to a timed phylogeny using VirusTreeSimulator (Ratmann et al. 2017), and that, in turn, was transformed to a tree with branch lengths in units of substitutions per site by simply assuming a strict molecular clock with a substitution rate of 0.001 substitutions per site per year (i.e. by multiplying each branch length by 0.001). This was then used to generate sequences with AliSim, with 10,000 bases, the GTR substitution model with equal base frequencies, and the continuous gamma distribution (shape = 0.5, scale = 2) as the rate heterogeneity model. We did this 50 times to get 50 alignments. For each of the 50, we picked 500 random sequences and progressively downsampled these to 100 in decrements of 100. We then performed that downsampling procedure 4 more times each. This gave us a total of 1250 alignments, 250 of each size. We ran IQ-TREE using both the DGM and FreeRate on all of them, and used its empirical Bayes routine to estimate site-specific substitution rates.

In figure 8, left, the x-axis represents the ratio of the sum of all branch lengths of the reconstructed DGM tree to that of the true tree. The y-axis is the reciprocal of 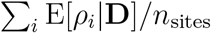 estimated by empirical Bayes. Grey lines join results from the same sequencing and down-sampling replicate. There is a very clear correlation, albeit with some noise around the line of equality. However, it should be noted that the maximum likelihood estimate of a phylogeny derived from a simulated alignment would not be expected to exactly match the tree used to generate that alignment. We can, however, get an estimate for the ML phylogeny of the simulated alignment by using FreeRate (Fig 8, right). Here the points are very close indeed to the line of equality, albeit with increasing deviation from it as tree sizes get smaller. We are unsure about the reasons for the latter phenomenon, which may simply be to do with limitations of the FreeRate model or the heuristic maximum likelihood approximation, and leave it for future work.

**Figure 8:**
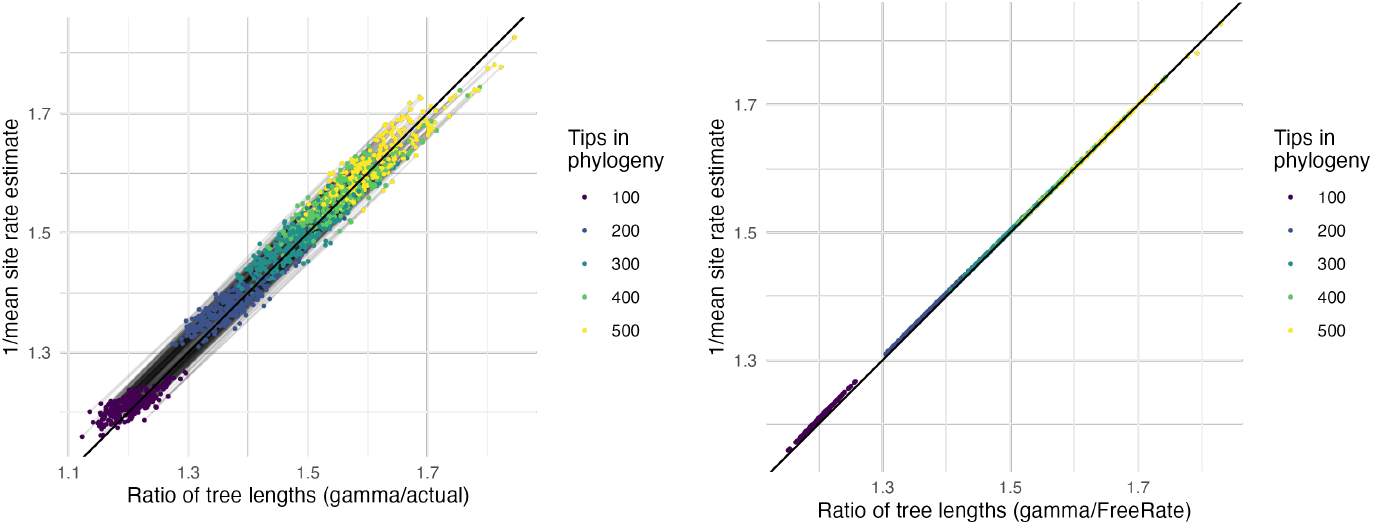
Left: For 1250 simulated alignments (points), we plot, on the x-axis, the ratio of the total tree length of a phylogeny reconstructed using the DGM to the total length of the true tree, versus, on the y-axis, the reciprocal of mean reconstructed site rate across the genome. The thick black line is equality; thin black lines connect alignments from the same simulation and downsampling replicate. Right: The same, except the x-axis is now the ratio of the total branch length from a DGM reconstruction to the total branch length from a FreeRate reconstruction.

As branch lengths are a measure of the amount of elapsed mutation, it is entirely to be expected that if per-site substitution rates are systematically mis-estimated then lengths would also be. The true *ρ*_*i*_s have a mean of 1 by definition, but their estimated values may not. In the next section we show this formally for an equal-rates model in our simplified scenario of a star-like tree.

### Likelihood in the large *n*_tips_ limit

To recap, *ρ*_*i*_ is the genuine relative substitution rate at site *i*, drawn from the distribution *H*. The mean *ρ*_*i*_ over **D** is 1 and thus the *ρ*_*i*_s sum to *n*_sites_. The probability of seeing a substitution at *i* on any branch is 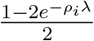 . In contrast, 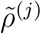is one of the *n* _cats_ discrete rates in the DGM which each *D*_*i*_’s rate equals with probability 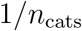. Define 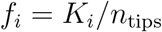. Now, at a single site *D*_*i*_:

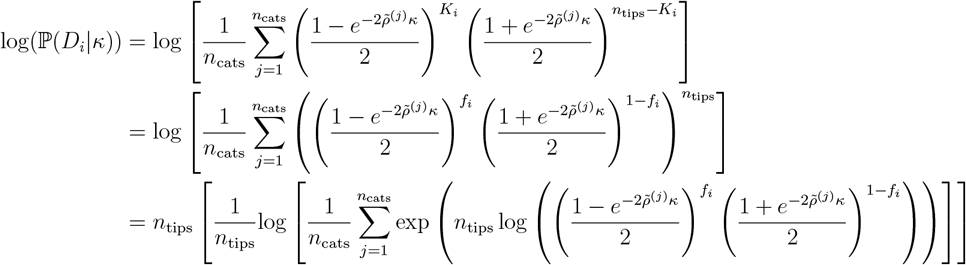

As is often the case with log-likelihoods, this analytical form is related to the log-sum-exp family of functions, part of the “softmax” functions widely used in machine learning. In fact, the last line is *n*_tips_ times the “mellowmax” function applied to 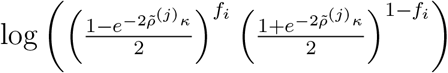, and this converges to the largest term in the sum as *n*_tips_ *→* ∞ (Asadi and Littman 2017). Note that, since the rates 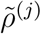are distinct, all terms in the sum in the above will be different and there will be a single category with a largest rate. Therefore we can approximate the above as

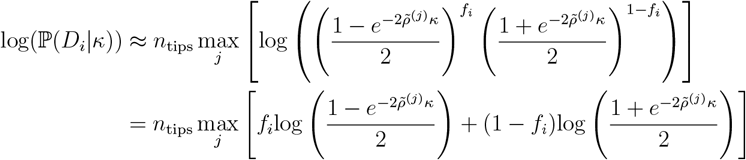

and the log-likelihood (6) can be written (up to additive constants) as

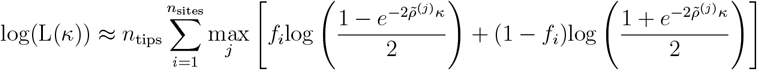

The value of *f*_*i*_ is fixed by the data rather than dependent on the value of *κ*, and for large *n*_tips_, it converges towards its expected value 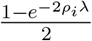. Therefore, the log likelihood can be well approximated by

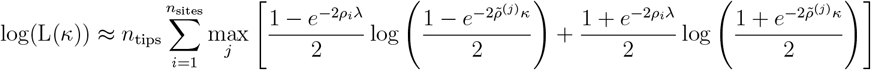

Suppose *λ* is sufficiently small that we can approximate 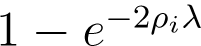 as 2*ρ*_*i*_*λ*. If the value of *κ* which maximises the likelihood is not similarly small, then the existence of considerable bias in the reconstruction of branch lengths is trivial, so we also confine our possible values of *κ* to those such that 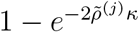can be approximated as 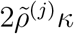. This leads to

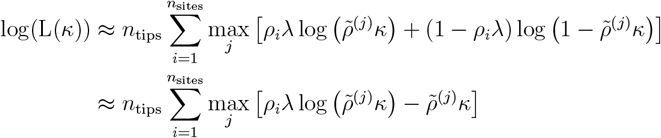

where in the last expression we ignore quadratic and higher terms in *λ* and *κ*. In order to move the maximisation over *j* out of the summation, separate it into a variable *j*_*i*_ for each position *i*:

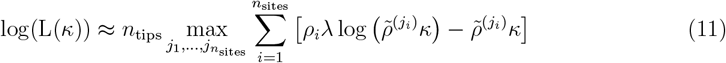

Note that if 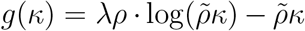then 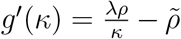and *g* is maximised when 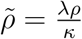, and thus if *λ* = *κ* and the 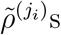can be freely assigned to any value (i.e. *n*_cats_ = *n*_sites_; a FreeRate model with one category per site), the maximum likelihood is obtained when each 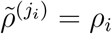.

We now want to find 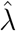, the value of *κ* that maximises (11). That maximum is approximately:

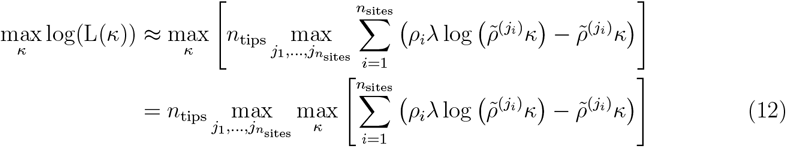

The first derivative of the internal expression with respect to *κ* is

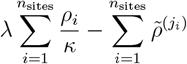

which gives a maximum at:

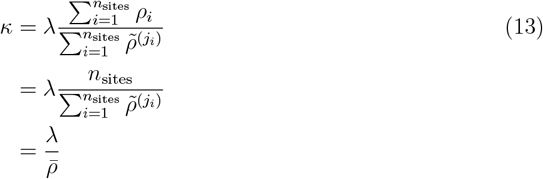

where 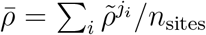. This is the value of *κ* that maximises the likelihood, and the value of that likelihood is

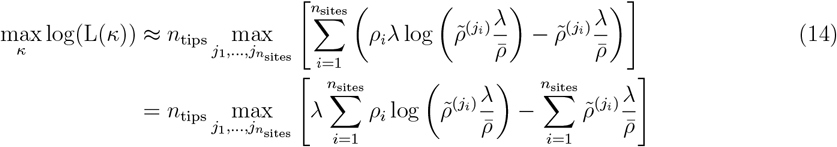

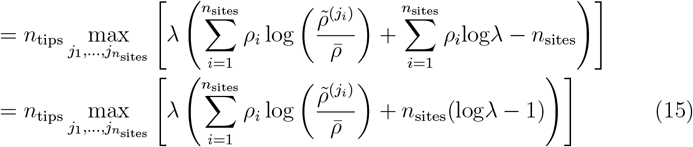

Every site *i* has a value of *j*_*i*_, 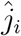, that maximises 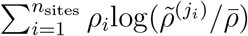. Maximising all of them simultaneously gives the maximum likelihood estimate of *λ* as

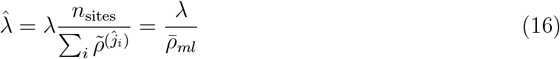

where 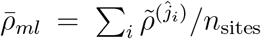. This formally demonstrates the validity of the findings of the previous section for our simplified tree structure, showing how the maximum likelihood branch length under an equal-weights model will not necessarily equal the true one. In the next two sections, we go on to show how explicit mismatches between the true generating distribution and that of the reconstruction method can cause the phenomenon we describe in this paper. We first consider a discrete generating distribution, and secondly a continuous one.

### And equal-weights model misapplied to a discrete generating model with unequal rates

If (11) is the approximate log likelihood for a given *κ* integrating over site rate assignments, then

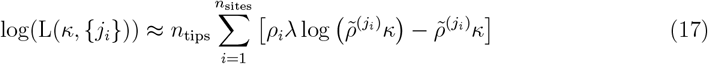

is the approximate log likelihood for a given *κ* and known rate assignments.

Consider the case where the true site-specific rates are defined by a FreeRate distribution with two kinds of sites, with “slow” and “fast” rates *ρ*^*s*^ and *ρ*^*f*^, *ρ*^*s*^ < *ρ*^*f*^ . The weights are *w*^*s*^ and *w*^*f*^ such that *w*^*s*^ + *w*^*f*^ = 1 and *w*^*s*^*ρ*^*s*^ + *w*^*f*^ *ρ*^*f*^ = 1.

In the last section we showed that if each site is assigned to its correct underlying rate and *λ* = *κ* then the likelihood is maximised when each *D*_*i*_ is assigned its correct rate. It should also be clear that if each 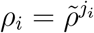then *λ* = *κ* in (13). Thus the maximum likelihood values of *λ* and the 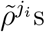are the ones that generated the data. The log value of this likelihood is approximately:

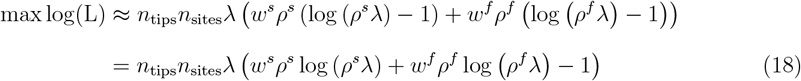

Now consider a model with rates 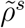and 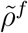such that 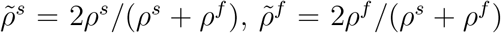and equal weights of 1/2. (Equal weights are actually required for the mean rate across sites to be 1 here.) If we assign 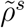as the rate to the *n*_sites_*w*^*s*^ “slow” sites and 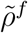to the *n*_sites_*w*^*f*^ “fast” ones, the mean rate across sites is:

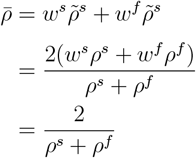

The likelihood for this rate assignment and a branch length of 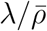is:

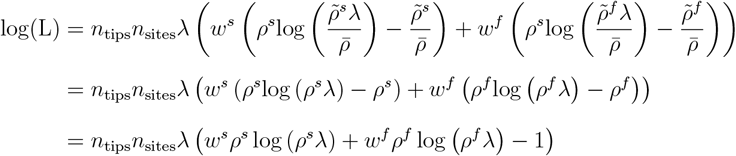

This matches (18). Thus the likelihood of the equal weights model with rates 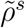and 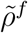 is equal to the maximum likelihood of the FreeRate model. As the equal weights model is nested in the FreeRate model, this must also be the *maximum* likelihood of the former.

So the likelihood of the equal weights model is maximised by assigning individual rates to sites in a way that departs from the equal weights assumption. But when this is done, the mean rate 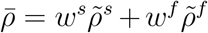is not 1 and thus the maximum likelihood estimate 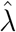of the branch length is not *λ*.

The relative bias in the branch length estimate is given by 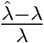and by using the identities 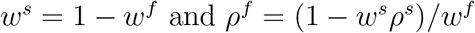this can be written as:

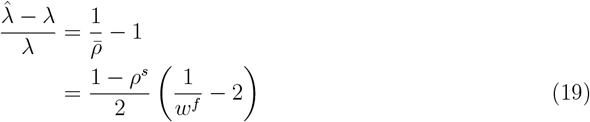

This result shows that a) the magnitude of the bias is larger for a larger separation between fast and slow rates (bias increases as *ρ*^*s*^ *→* 0) and for a smaller fraction of rapidly evolving sites (bias increases as *w*^*f*^ *→* 0). As *ρ*^*s*^ is always less than 1, the sign of the bias is driven by *w*^*f*^ : if this is greater than 0.5 then branches will contract; otherwise they expand. The former case is rather unlikely in real-world situations, at least in the biological realm, where rate distributions generally have a mode of less than 1 (Jia, Lo, and Ho 2014; Silvestro, Latrille, and Salamin 2024).

To demonstrate this, we repeated the first simulation exercise of the main text (“A bias in reconstructed branch lengths exists for the DGM but is absent for FreeRate”), for the same set of trees, with the generating rate heterogeneity model replaced by a four-category FreeRate model with rates 0.1, 0.4, 0.9 and 1.6 and corresponding weights 0.1, 0.2, 0.3 and 0.4. For this instance of a positive correlation between weights and rates, the result is a branch contraction effect when the DGM is used for reconstruction (figure 9).

**Figure 9:**
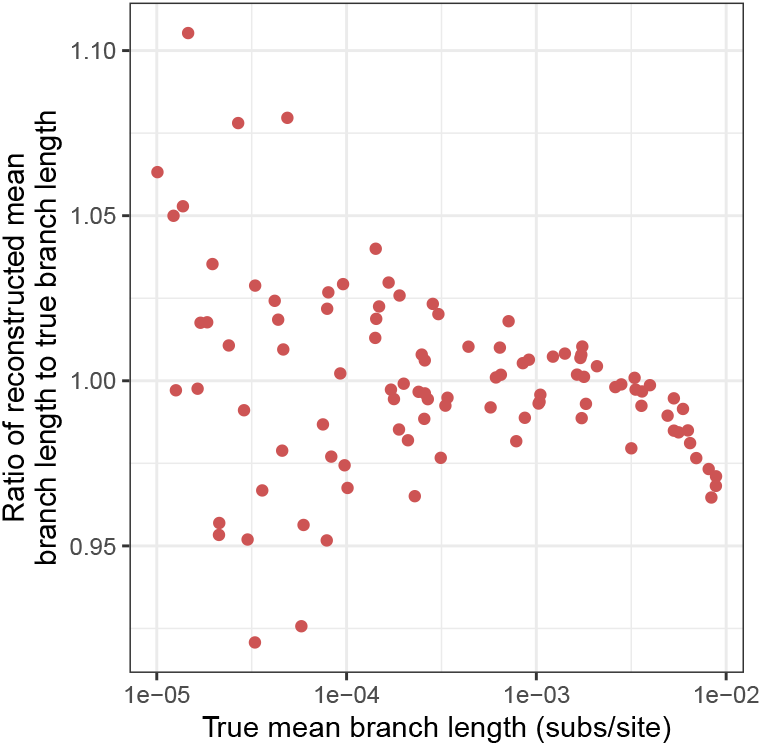
For 100 simulated phylogenies with a variety of mean branch lengths, the ratio of reconstructed to true mean length plotted against the true mean, when sequences were simulated using a 4-category FreeRate model with rates and weights positively correlated, and the reconstruction model is the 4-category DGM.

We can generalise this to the situation of a tree generated using a *n*_cats_-category FreeRate distribution and reconstructed with an equal-weight model also with *n*_cats_ categories. Denote the rates and weights of the FreeRate distribution by *n*_cats_-dimensional vectors **P** = {*ρ*^(*c*)^} and **W** = {*w*^(*c*)^} satisfying 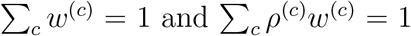. If we assign site-specific rates as 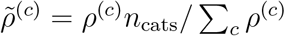and assign the sites to categories correspondingly, these rates correspond to the maximum of the unconstrained likelihood (13), and thus must maximise the likelihood of the reconstruction model. The value off 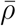or this realisation can be easily computed as 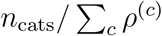and therefore 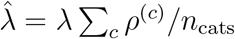. Then the relative bias can be written as:

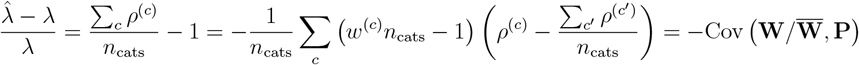

Thus the bias is inversely proportional to the covariance between rates and relative weights among categories. When the distribution includes a tail of rare but fast-evolving sites, there is a negative correlation between weight and rate and therefore a branch expansion effect.

### An equal-weights discrete model as an approximation for a continuous Gamma generating model

Here we allow the underlying rates to be drawn from the simplest possible continuous Gamma distribution, that with shape and scale 1 (i.e. an exponential distribution with rate 1). We will fit a two-category discrete model with equal weights. As in the previous section, we take two rates 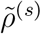 and 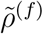, with 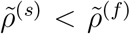, such that 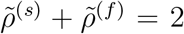. We can write *ρ*^(*s*)^ = 1 − *ϵ* and *ρ*^(*f*)^ = 1 + *ϵ* for 0 *≤ ϵ ≤* 1. Both weights are 1/2. In contrast to the previous sections, where category rates are known, we do not assume we know *ϵ* and must estimate it.

The expression which is to be maximised with respect to both the rates and their site assignments is derived from (23):

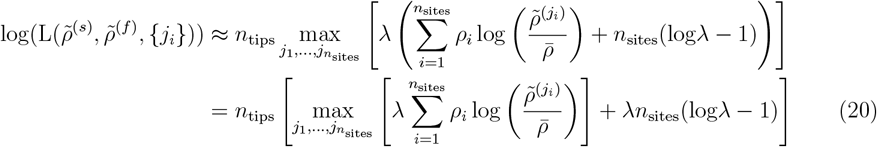

where each *j*_*i*_ ∈ {*s, f*}. We first need to show that, for some *α* between 0 and 1, the likelihood is maximised when the slowest *αn*_sites_ sites are allocated to 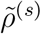and the fastest (1 − *α*)*n*_sites_ to 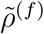. To see this, suppose you know the value of *α* that maximises the likelihood L and calculate L under this allocation. Now switch the assignments of two sites *a* and *b*, where *a* was assigned to 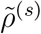and *b* to 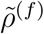in the assignment maximimising L, and recalculate the likelihood as L^*′*^. Using (14) as the expression for the likelihoods, L and L^*′*^ differ only in as much as L contains the term:

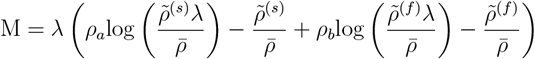

while L^*′*^ contains:

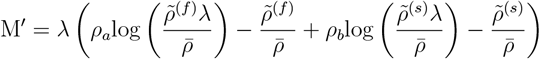

where 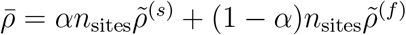.It remains solely to show that

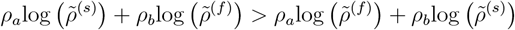

but straightforwardly:

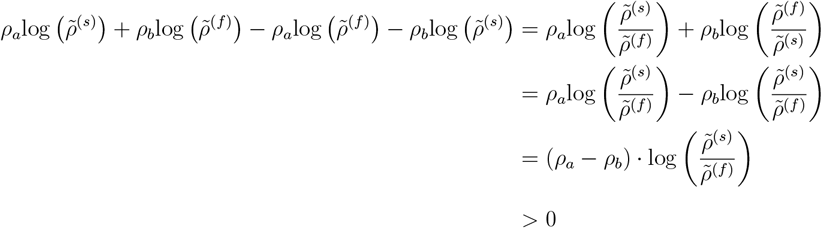

(the last because both 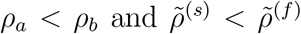. Thus the maximum likelihood allocation of sites to rates must work in this way and it remains only to estimate *α* and *ϵ*.

Rearrange the index of the sum in (20) such that *ρ*_*j*_ *> ρ*_*i*_ if *j > i*. To get an analytic expression for the log likelihood, consider that

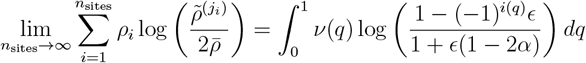

where *ν*(*q*) = − log(1 − *q*) is the quantile function of the true rate distribution and *i*(*q*) = 1 if *q ≤ α* and 0 otherwise. If we take *n*_sites_ to be large enough that the integral can be taken as an approximation of the sum, then the term *λn*_tips_*n*_sites_(log *λ* − 1) in (20) is invariant by site and it remains only to maximise the integral over *α* and *ϵ*

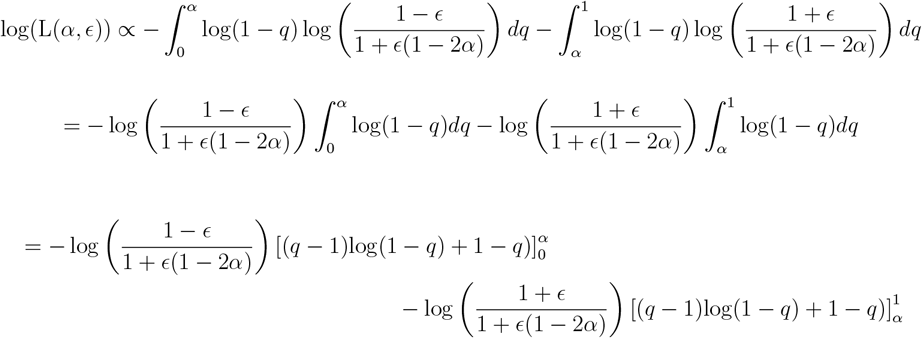

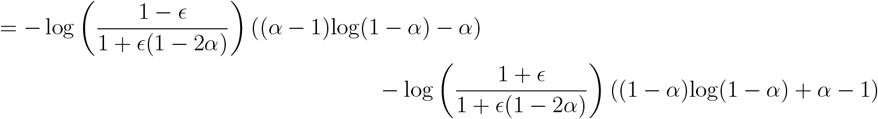

To maximise this, we calculate its partial derivatives with respect to *ϵ* and *α*, to obtain:

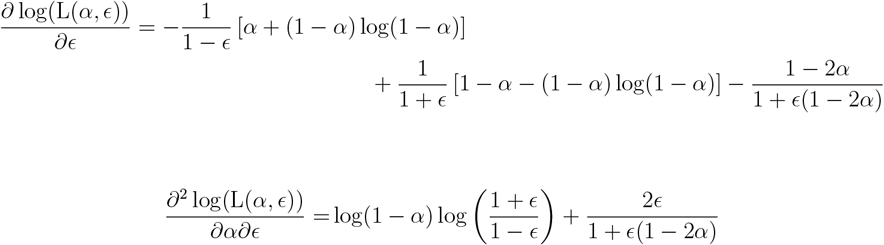

If solved numerically this gives *ϵ ≈* 0.65 and *α ≈* 0.64, from which 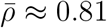. If we look at the expression for of 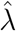 from equation (24), the maximisation of the likelihood results in a branch expansion of roughly 20%.

### Generalisations

In this section, we show how some of the assumptions used in the mathematical derivation above can be relaxed without affecting the main results.

#### Unequal branch lengths

While keeping the star tree, assume that each branch has a different length *λ*_*a*_. We assume that these lengths are bounded and that the sum of all lengths diverges for large *n*_tips_. The likelihood for the entire alignment becomes

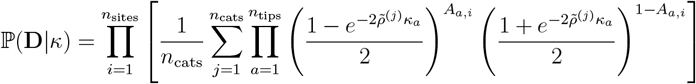

where *A*_*a,i*_ represents the allelic state of tip *a* at site *i* in the multiple sequence alignment. In the limit of large *n*_tips_, we obtain (up to a constant)

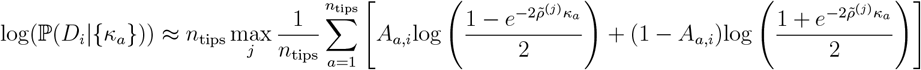

where the *κ*_*a*_ are the variables representing reconstructed branch lengths. Each *A*_*a,i*_ is a Bernoulli random variable. It has mean 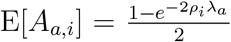 and variance 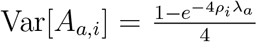 The function of *A*_*a,i*_ inside the square bracket is a random variable *B*_*a,i*_ with:

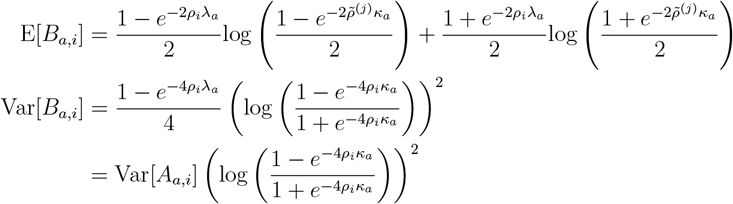

We want to be able to use the Lyapunov Central Limit Theorem (LCLT), in which case we need to show that there exists *δ >* 0 such that:

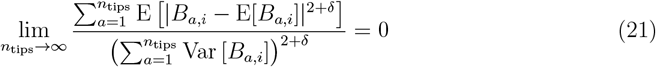

As 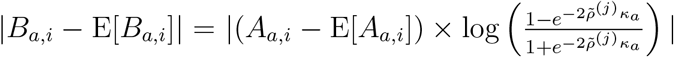, and 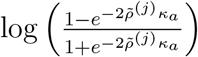is negative and independent of *A*_*a,i*_, this becomes:

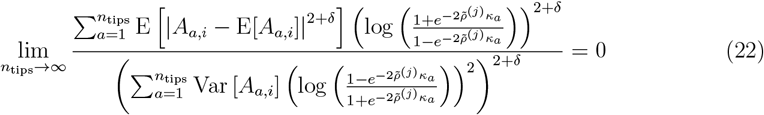

This is satisfied for *δ >* 0 provided that both *λ*_*a*_ and *κ*_*a*_ have a finite non-zero upper and lower bound, since in this case all terms in the numerator are bounded from above by 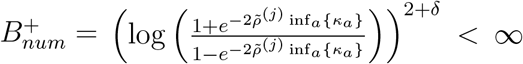 and all terms in the denominator are bounded from below by 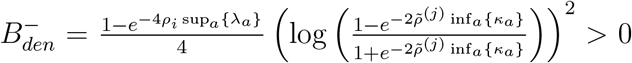, and the denominator scales as 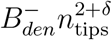 while its numerator scales as 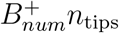. This condition also implies that the total tree length scales linearly with the number of tips *n*_tips_.

Therefore, under the condition that branch lengths are bounded (and hence the means and variances of the *B*_*a,i*_s are finite), the LCLT implies that the sum in the expression for the likelihood above converges to its mean value as *n*_tips_ *→* ∞, resulting in the likelihood (using the same assumptions of small *λ* and *κ* as before)

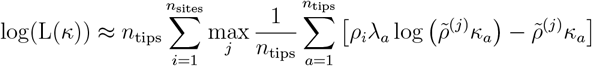

which is maximised by 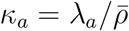,Substituting this, we end up with the likelihood maximisation

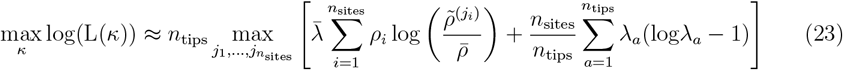

which is the same as the original expression (23) up to a constant that is irrelevant for the maximisation, and up to the replacement of *λ* (identical for all branches) with the average branch length 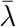. Therefore, the values 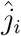 of *j*_*i*_ that maximise this expression do not change, and the maximum likelihood estimate of *λ*_*a*_ becomes

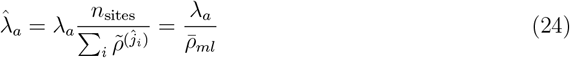

so the ML estimates for all branches are the same multiple of the true value as before (“Likelihood in the large *n*_tips_ limit”).

#### Asymmetric substitution model

For an asymmetric substitution model, assume two states *A* and *B* with true substitution rates *µ*_*A*_ and *µ*_*B*_ with the constraint 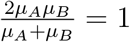. The equilibrium probabilities for the two states are 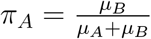 and 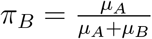. We assume that the root sequence is a mixture of the two states with equilibrium proportions. The unknown versions of these parameters to be inferred in the likelihood are indicated by 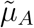and 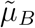.

For site *i*, assuming that the root state is *A* and the count of tips with *B* is *K*_*i*_, the likelihood is

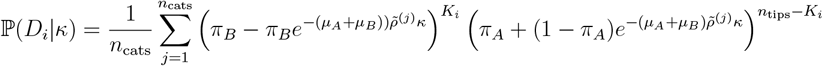

With the same approximations as before, this becomes

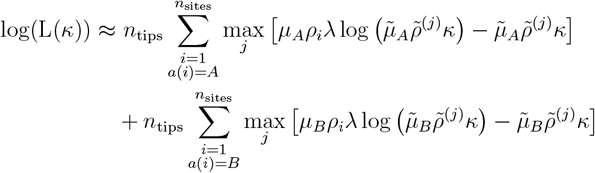

Assuming no correlation between the root state and the site-specific rates, this likelihood is maximised when 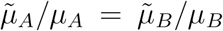, which implies 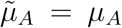 and 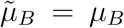so as to satisfy 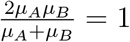 The likelihood is then reduced to

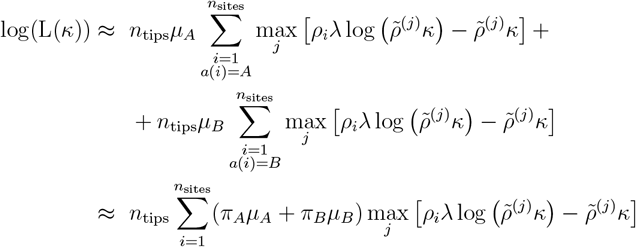

which reduces to the usual form for the likelihood since *π*_*A*_*µ*_*A*_ + *π*_*B*_*µ*_*B*_ = 1, and in turn leads to the same maximum likelihood estimates of branch length.

#### Unknown root state

If we assume that the root state is unknown, all likelihoods in the previous sections should have an additional term that represent the probability of the phylogeny if the root state is 1. After usual approximations, the likelihood becomes

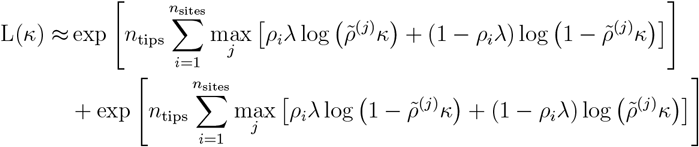

and the relative contribution of the second term to the likelihood is negligible, since it is bounded by

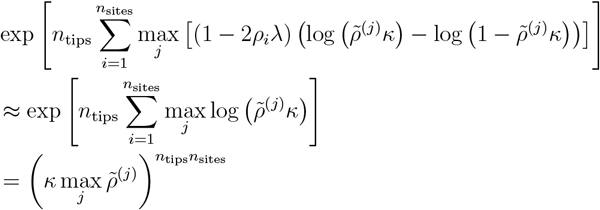

which tends to zero as tends to infinity. Therefore, the contribution of the alternative root state can be ignored and all results are unchanged.

### Open questions

There is scope for further unpacking the exact cause and properties of this bias. In particular, in this work we do not demonstrate the directional effect of adding more tips to the tree, instead concentrating on the large *n*_tips_ limit. As the magnitude of the effect of increasing sample size eventually levels off (figures 2, 7), the biases calculated in this appendix presumably correspond to the asymtomatic limit, but a rigorous demonstration of this would require further work. Related to this the observation from simulations that rate at which the bias increases is correlated with average branch length (figure 7); this is the likely reason that the problem is often not discernable when the latter are small.

