## Supplemental figures for "Biased estimates of phylogenetic branch lengths resulting from the discretised Gamma model of site rate heterogeneity"

September 23, 2025

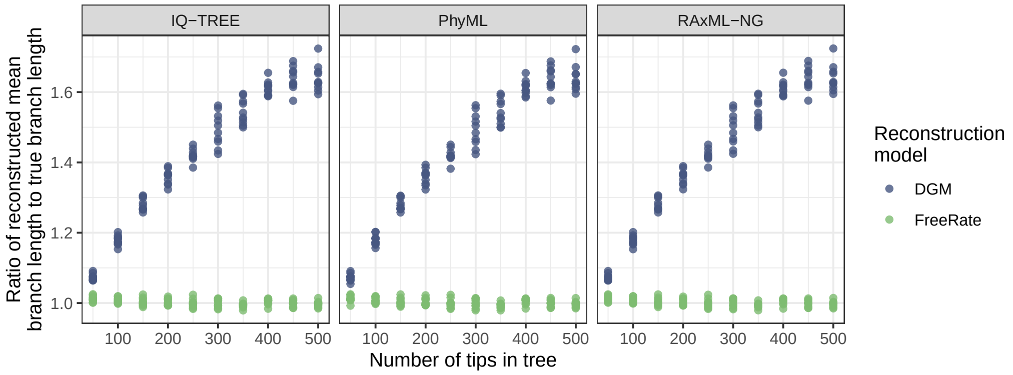


Figure S1: The effect of the number of tips in the tree, as reconstructed by different maximum likelihood packages. For a total of 100 simulated trees with varying numbers of taxa, the ratio of the reconstructed mean branch length to the true mean branch length (which was always 0.01 substitutions per site) is plotted. The generating rate heterogeneity distribution was a continuous gamma distribution; the reconstruction was done with the DGM (blue) and FreeRate (green). The facets are the software package used to for reconstruction.

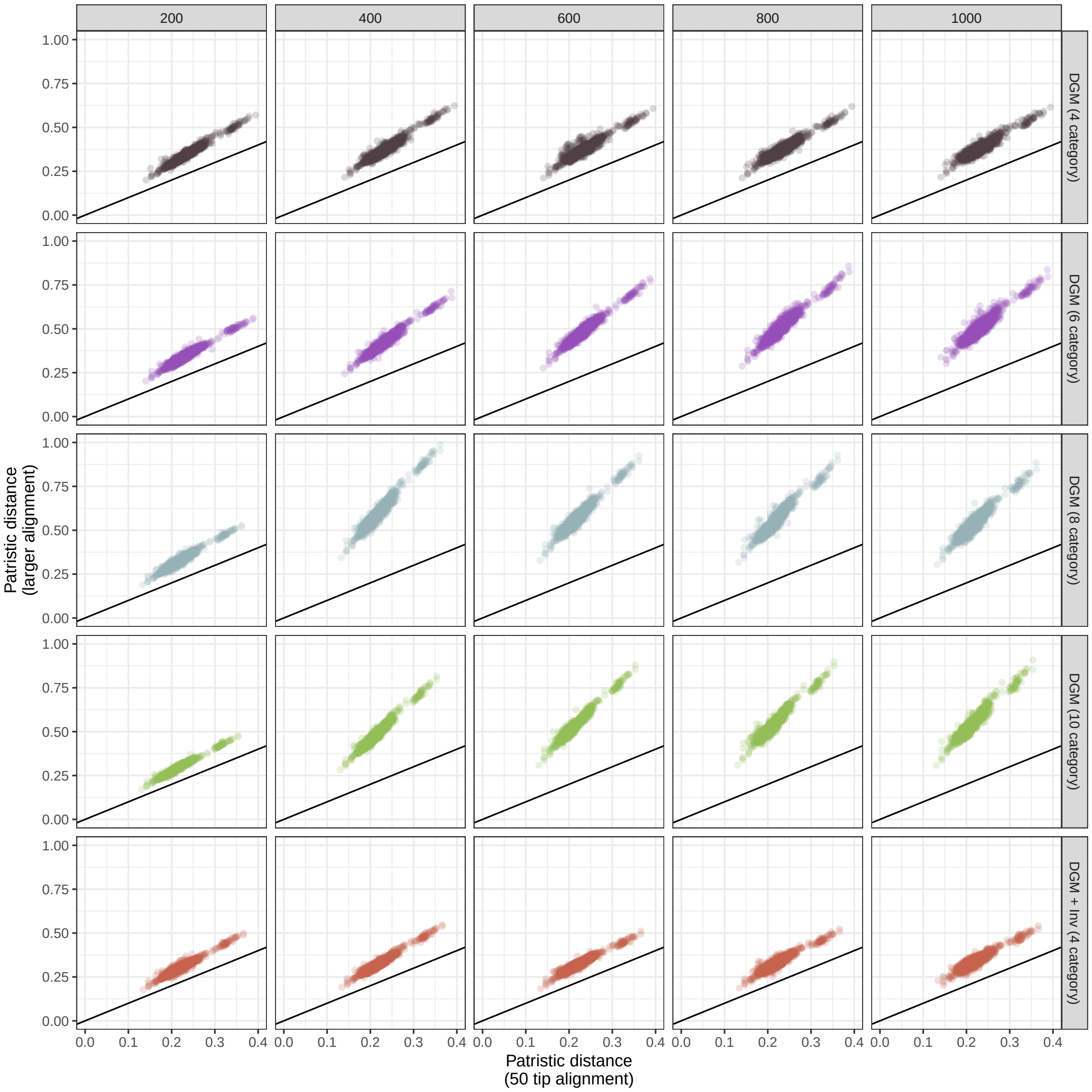


Figure S2: Patristic distances between the tips representing 50 subtype C HIV sequences in phylogenetic reconstructions. The x-axis gives these when the alignment consists of just those 50 sequences, while the y-axis are the same from a larger alignment, with the total number of sequences increasing from left to right in the horizonal facets. Facet titles are the sizes of these larger alignments. Reconstruction was done with a variety of different versions of the DGM (vertical facets).
